# The archives are half-empty: a field-wide assessment of the availability of microbial community sequencing data

**DOI:** 10.1101/2020.04.28.063271

**Authors:** Stephanie D. Jurburg, Maximilian Konzack, Nico Eisenhauer, Anna Heintz-Buschart

**Affiliations:** German Centre for Integrative Biodiversity Research (iDiv) Halle-Jena-Leipzig, Deutscher Platz 5e, 04103, Leipzig, Germany; Martin Luther University Halle-Wittenberg; Leipzig University, Institute of Biology, Deutscher Platz 5e, 04103 Leipzig, Germany; Helmholtz Centre for Environmental Research GmbH - UFZ

**Keywords:** data availability, microbiome research, meta-analysis

## Abstract

The sequencing revolution has resulted in the explosive growth of public genetic repositories. These repositories now hold invaluable collections of 16S rRNA gene amplicon sequences, but the extent to which the currently archived data is findable, accessible, and reusable has not been evaluated. We conducted a field-wide assessment of the availability and state of publicly archived 16S rRNA gene amplicon sequencing data. Using custom-built pattern-based text extraction algorithms, we searched 26,927 publications in 17 microbiology or microbial ecology journals, and identified 2,015 studies which performed 16S rRNA gene amplicon sequencing. We found, for example, that 7.2% of these had not been made public at the time of analysis, a trend which increased over time. Of the 635 studies targeting the V3-V4 region of the 16S rRNA gene, 40.3% contained data which was not available or not reusable, and for 25.5% of the studies, faults in data formatting or data labelling were likely to create obstacles in data reuse. Taken together, only 34% of these datasets had potentially reusable data. Our study reveals significant gaps in the availability of currently deposited community sequencing data, identifies major contributors to data loss, and offers suggestions for improving data archiving practices in the future.

## Introduction

Advances in microbiological research have been marked by the steady development and optimization of sequencing technologies. Where culture-dependent methods forced microbiologists to focus on a small portion of the world’s microbes (1, 2), high throughput sequencing methods allowed the field to bypass this limitation and indirectly observe complete microbiomes at increasingly high resolutions. As a result, the last decade has also seen an exponential growth in the number of studies producing sequencing data, as well as in the quality of this data (3).

In particular, 16S rRNA gene sequencing, by which a section of the small ribosomal subunit’s RNA gene is amplified, sequenced, and used as a tag to identify prokaryotic taxa (archaea and bacteria), has prospered during this time. 16S rRNA gene amplicon sequencing has provided microbial ecologists with census data for microbial communities similar to, or often more complete than, those obtained by macroecologists during field sampling, for example. High-throughput sequencing has supported research into the ecology of microbial communities, and a renewed interest in microbiome research. To date, individual studies have found parallels between the ecological patterns of microbiomes and those found for macroecosystems e.g., ecological scaling, species-abundance distributions, species-area relationships, and distance-decay (4, 5). However, the generalizability of these findings across ecosystems requires the systematic meta-analysis of ecological patterns across microbial communities in different environments (5–11)

Importantly, the uniform format of sequencing data has favored archiving practices (12, 13), and the fields of genetics and molecular ecology are often cited as pioneers within ecology (14). As microbial ecology has moved towards an increasing reliance on sequencing, the deposition of the resulting sequencing data into public genetic databases has become standard practice, and often a prerequisite for the publication in peer reviewed journals. Presently, the archiving of sequencing data is centralized in public genetic repositories belonging to the International Nucleotide Sequence Database Collaboration (INSDC): NCBI’s Sequence Read Archive (SRA), the EBI’s European Nucleotide Archive (ENA), and DDJ’s Sequence Read Archives (DRA) (15). These are regularly synchronized and support compatible data format, creating an opportunity for synthesis in microbiome research. Recent meta-analyses of publicly available sequencing data have advanced the fields of medicine (9, 16), microbiology (10), and microbial ecology (11). It is expected that future advances in these areas will rely heavily on sequences which have been archived (5, 7); however, the degree to which the data which is currently archived is reusable in syntheses has not been evaluated.

For archived data to serve synthesis efforts, they must be stored in findable, accessible, interoperable, and reusable formats (12). Accordingly, INSDC databases require users to provide experiment and sample-level metadata (17), in addition to raw sequence data, and in turn provide the users with stable accession numbers and the long-term storage of their data. To evaluate how much of the currently deposited sequencing data may serve as a resource for future syntheses, we performed the first comprehensive assessment of data availability and reusability in microbial ecology. We surveyed all literature in the 17 most prominent microbial ecology-related journals since 2015 (n= 26,927 articles), and using a custom-built pattern-based text extraction algorithm followed by manual curation, selected those studies which performed 16S rRNA gene amplicon sequencing and listed INSDC-compliant accession numbers (n= 2,015, Table S1; 145,203 samples) to assess whether the data was available, accurately labeled, and properly formatted to serve future synthesis efforts.

## Materials and Methods

### Journal selection

An initial survey of the literature was performed in February 2019 on Google Scholar with the following search query: *“bacteria” AND “515” AND “806”* to obtain a preliminary assessment of the literature employing amplicon sequencing V3-V4 hypervariable region of the 16S rRNA gene, which has been recommended and popularized by the Earth Microbiome Project (18). Results were filtered to include only publications since 2015 and yielded ∼8,600 hits. The software Publish or Perish (19) was used to obtain general bibliographic information for the first 1,000 hits for the Google Scholar query (refined to *bacteria AND 515 AND 806 NOT book patent*) for each year between 2015 and 2019, yielding 4,635 results (available in Supplementary Table S1). The 17 most common journals in this list were considered the main publishers of microbial ecology data (Table S1) and were selected for further analyses. The preprint server bioRxiv was excluded, because we only considered work that had passed the reviewing process. Similarly, we excluded journals which were not specialized in microbiology or microbiome research (i.e., *PeerJ, Nature Communications, PLOS ONE*), as specialist journals had more specific and stringent requirements for data deposition, and the authors of articles in these journals were more likely to be acquainted with microbiome data.

In March 2019, all articles from each of the selected journals published between January 2015 and March 2019 were downloaded as follows: the DOIs for all publications for each journal for the period studied were obtained by querying the Web of Science, using the Publish or Perish software. The concatenated list of DOIs was entered into Citavi (https://www.citavi.com) to create a bibliography and to download the corresponding articles as PDFs. Articles for which the PDFs could not be downloaded were excluded. The final set included 26,927 articles.

### PDF preprocessing, text mining, and article selection

After renaming the entire corpus, we checked the PDF format for each file using *pdfinfo* of poppler tools (https://poppler.freedesktop.org/). We excluded invalid pdfs (n=12), and applied the command *pdftotext* to extract plain text from each pdf. For each article, a corresponding searchable TEI XML file (https://tei-c.org/) was created using the GROBID v.0.5.4 command *processFulltextDocument* (https://github.com/kermitt2/grobid/). In 218 cases, GROBID was not able to generate such XML documents, and these were excluded from further analyses. For the extraction and parsing of each TEI document, we developed and implemented a customized python package (https://github.com/komax/teitocsv). Briefly, each TEI document was parsed and searched for occurrences of general patterns including author, DOI, and journal. From the title, abstract, and main text fields (excluding references and supplementary materials), we extracted patterns indicating their relevance to this study, in particular accession numbers corresponding to INSDC-associated databases (specifically PRJ, ERP, DRP, SRP, SAME, SAMND, SAMN, ERS, ERX, DRX, SRX, DRR, SRR, ERZ, DRZ, SRZ followed by six digits) and references to the 16S rRNA gene, high throughput sequencing platforms, and the 16S rRNA gene region sequenced (i.e., primers). These data were outputted as a single CSV file summarizing the findings for the entire corpus, with each accession number occupying a separate row (multiple rows per article possible) and each column capturing an aspect of pattern matching (i.e., DOI, sequencing platform). A detailed flowchart of the article selection process is included in Supplementary Figure S7. The accuracy of our parsing methodology was confirmed by manually inspecting 150 randomly-selected articles which mentioned 16S but for which no accession number or alternative database was detected (Supplementary Data). Of these, only one article contained an accession number which was incorrectly reported (i.e., missing characters), and two had deposited their data in unconventional locations (google drive and the GEO database). Of the 150 articles inspected, none had deposited sequence files or accession numbers in the supplementary section. These articles were also used to estimate the number of articles which described 16S rRNA amplicon sequences but did not provide an accession number for the stored data.

To assess the completeness of our data relative to all available amplicon sequencing datasets currently in existence, we conducted a Web of Science search for all articles citing the Mothur (20) or QIIME (21, 22) bioinformatic tools for processing amplicon sequences on March 10, 2020, excluding all publications which were not articles or early-view articles, and had been published between 2015-2019. These workflows are the most common tools for processing amplicon sequences, hence either one is likely to be cited in articles reporting 16S rRNA gene amplicon sequencing data. Among our 17 target journals, they were cited by 1,984 articles. To access the sequencing data, ranges of accession numbers within articles were resolved to single accessions in a two-step process: first, all potential ranges were defined based on the occurrence of multiple accession numbers with the same prefix within the same manuscript. Secondly, ranges larger than 40 accessions per study were verified manually and smaller ranges were included automatically, because all ranges between 30 and 40 were found to contain true ranges of accession numbers in a manual check. False positive accessions introduced in this step were manually removed during the final analysis.

### Data access

Run-level metadata in the sequence read archive were mined for all accessions via the NCBI’s Entrez Direct (EDirect) toolkit’s *esearch* and *efetch* (esearch -db sra -q <u><ACCESSION</u>> | efetch - format runinfo) on the 28^th^ June 2019. We manually curated 964 articles containing accession numbers that did not yield any data, or that yielded metadata which was not labelled with the library strategy “AMPLICON” to exclude 577 articles from further analysis that did not report amplicon sequencing results of phylogenetic marker genes. Accession numbers from articles that were verified to report amplicon sequencing results but did not lead to sequence read sequencing metadata were manually confirmed on the NCBI web portal.

To assess the validity of the submitted (raw) sequencing data, the accessions were reduced to those mined from articles which mentioned the most frequently used primer pair 515F and 806R (8, 23) (Supplementary Figure S8). Specifically, for the 515F primer, we captured any combination of the occurrence of 515 F(wd) or F(wd) 515 which was separated an arbitrary number of white spaces in addition to barcoded versions of the original and the modified 515f primer. For the 806R primer, we exhausted all variations of this primer analogously. Our procedure also captures non-minimal mentions (i.e, 806Rb). For this dataset (referred to as 515F-806R-substudy), 1,000 reads per submitted fastq-file were downloaded on July 2^nd^ 2019 using NCBI’s prefetch, vdb-validate and fastq-dump tools (runs specified in the library layout field as single-end sequencing: prefetch <RUN-LEVEL ACCESSION> && vdb-validate <RUN-LEVEL ACCESSION> && fastq-dump -X 1000 <RUN-LEVEL ACCESSION>; runs specified as paried-end sequencing: prefetch <RUN-LEVEL ACCESSION> && vdb-validate <RUN-LEVEL ACCESSION> && fastq-dump -X 1000 --split-3 <RUN-LEVEL ACCESSION> (24).

In studies containing only one fastq file, potential barcodes were extracted by trimming reads starting from the 515F or 806R primer sequences using cutadapt version 1.18 (25). The number of reads containing potential barcodes of 4-30 base length adjacent to the primers, and the number of different barcodes per study were assessed.

### Sequencing data evaluation

Sequences were searched for both primers in the degenerate form (26, 27) and their reverse complements allowing for 20% mismatches and requiring 10 bases overlap. Read qualities were assessed using FastQC v0.11.3(28).

### Metadata access

All metadata connected to the samples of the 515F-806R-substudy was accessed on August 4^th^ 2019 via the biosample accession numbers from the run-level metadata using the NCBI’s Entrez Direct (EDirect) toolkit (esearch -db sra -query <BIOSAMPLE_ID> | elink -target biosample | efetch -format docsum | xtract -pattern DocumentSummary -element Attribute@attribute_name,Attribute) (29). The retrieved metadata was collated per study using R.

### Data analyses

All analyses and visualizations were performed in R 3.6.1. (30). We identified studies which had targeted the V3-V4 hypervariable region of the 16S rRNA gene between base pairs 515 and 806 by selecting articles which contained the all of the following keywords “16S rRNA”, “V3”, “V4”, “515”, and “806”. To test for changes in the percentage of datasets which fulfilled a particular condition over time, we used a chi-squared test for trend in proportions. For the metadata analyses, we focused on the metadata supplied for the V3-V4 subset of studies. To assess the environments studied in this subset, we looked at the frequency of different environments reported in the “ScientificName” field. To assess the informative potential of different metadata fields, we divided the fields into ‘mandatory’ if they were present in all datasets, and ‘popular optional’ if they were present in more than 25% of the studies. Note that whether a field is mandatory may change over time as INSDC deposition policies are improved. To determine the informative potential of each of these fields, we divided the number of samples in each study by the number of factor levels.

## Results and Discussion

To confirm that our parsing algorithm did not miss accession numbers in articles on 16S rRNA gene amplicon sequencing, we manually inspected articles in which 16S rRNA was mentioned, but accession numbers were not detected. Of the 150 randomly-selected articles inspected, one contained a misspelled accession number, two had archived their sequences in unconventional repositories (Google Drive and GEO, a gene expression database, Supplementary Data), and 19 were identified as having performed 16S rRNA gene amplicon sequencing but had not included any reference to the data. We found no evidence that accession numbers or sequence data were stored in supplementary materials. From this subset, we estimate that 18% of the studies in our database (n=469) performed 16S rRNA gene amplicon sequencing but did not provide access to the data (Figure 1a). We found that an additional 6.5% of the studies had deposited their data in the Qiita (31), MG-RAST (32), and figshare databases (n=14, n=134, and n=24 studies, respectively). Of the estimated 2,656 studies employing 16S rRNA gene amplicon sequencing, 75.9% deposited their data to an INSDC database in the period studied (Figure 1a).

**Figure 1.**
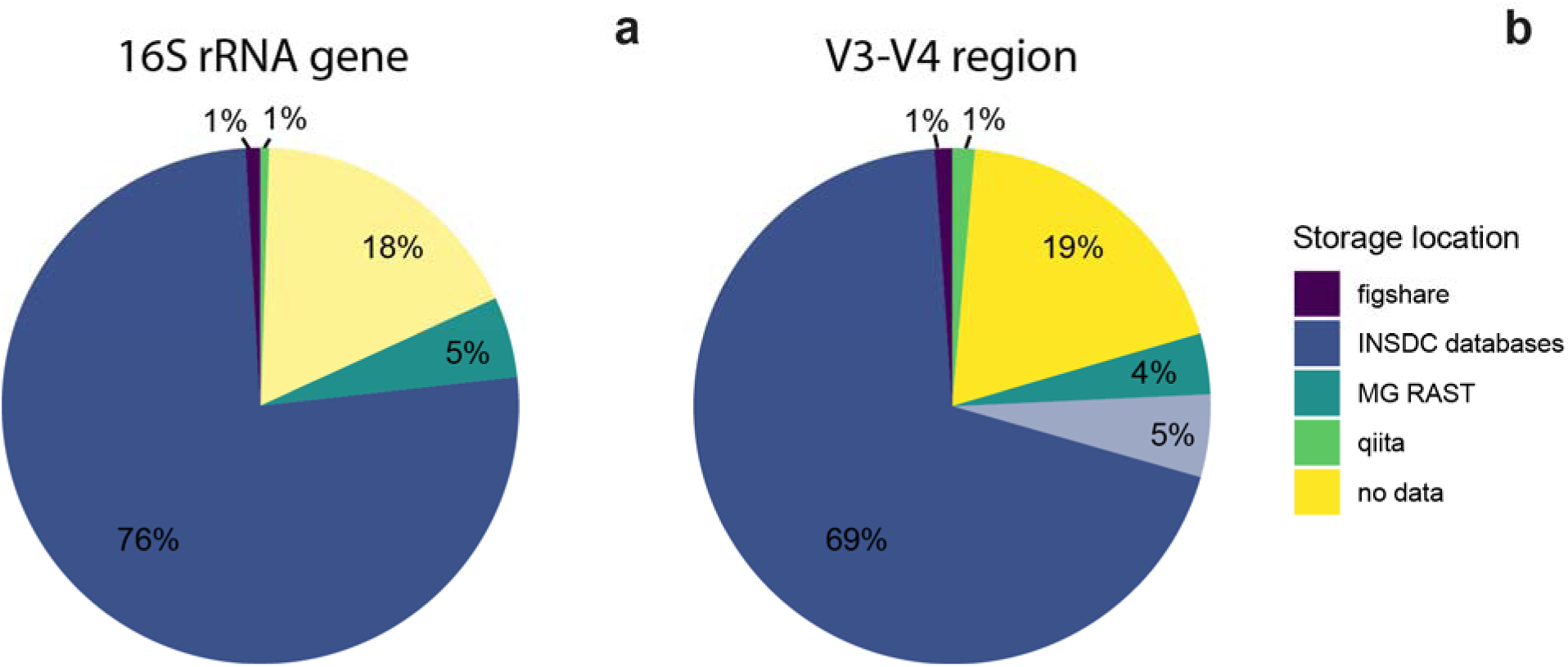
Popular locations for data storage. Data for all studies which contained 16S rRNA amplicon sequencing (a), and the subset of studies which targeted the V3-V4 hypervariable region of the 16S rRNA gene between base pairs 515 and 806 (b); n=2656 and n=635 studies, respectively. For the entirety of the study, studies which contained amplicon sequences but did not deposit them, were inferred by manually checking 150 randomly-selected articles which did not contain INSDC accession numbers or refer to alternative databases, indicated in lighter yellow. For the V3-V4 subset, studies which contained the keywords “16S rRNA”, “515”, and “806” were selected. Studies for which INSDC-compliant accession numbers were reported but which did not exist on any INSDC database are shown in lighter blue.

To obtain more precise estimates of the percentage of articles which deposited their data in each database, we focused on the subset of studies which sequenced the V3-V4 hypervariable region of the 16S rRNA gene between base pairs 515 and 806, a target region which has gained popularity since its development and use by the Earth Microbiome Project (18, 33). We identified the 635 qualifying articles by parsing our database for studies which contained the keywords “16S rRNA”, “515[f|F]”, and “806[r|R]”. Of these, 74.5% (n=474) studies listed INSDC-compliant accession numbers within the article, but of these, accession numbers from 33 studies were not findable on any INSDC database. Additionally, 19% (n=121) did not provide an identifiable link to the data, and 6.8% of the studies deposited their data in the Qiita, MG-RAST, and figshare databases (n=9, n=24, n=7, respectively, Figure 1b). Two studies provided SRA submission IDs rather than accession numbers, and were also inaccessible.

The increasing popularity of microbial community sequencing was evident in our data. Over the period studied, the number of studies targeting the V3-V4 region of the 16S rRNA gene rose from 56 in 2015 to 214 in 2018 (Supplementary Figure S1). The proportion of publications which claimed to deposit data to INSDC databases increased slightly over time, from 33/56 in 2015 to 172/214 in 2018 (χ^2^=6.6, *p*=0.01), suggesting an increasing tendency towards deposition in INSDC databases. Deposition to alternative databases decreased (χ^2^=14.04, *p*<0.001) indicating a switch to these standardized databases, but not towards making data accessible in general, as the proportion of studies which did not make their data public was remarkably stable over time (χ^2^<0.001, *p*=0.99). During this period, the number of studies without publicly available data rose, from 13 in 2015, to 38 in 2018 (Supplementary S1). Importantly, 5% of the studies (n=33) provided INSDC-like accession numbers, but these did not exist on any INSDC database, likely due to typos or not yet having been published.

While data deposition to any public repository is preferable over no deposition at all, non-INSDC alternatives were not designed for the long-term storage of 16S rRNA amplicon sequencing data, and thus are likely to lead to the long-term loss of information. Qiita’s intended use is “the analysis and administration of multi-omics datasets” (https://qiita.ucsd.edu/). This platform is not designed for the long-term archiving of these data, and accordingly, Qiita includes software to facilitate deposition of sequences to the ENA. Similarly, MG-RAST (32) is an online platform for metagenomics analyses which also facilitates sequence deposition to appropriate databases. In contrast, figshare is a general repository which hosts most forms of research output (https://figshare.com/), but it is neither sequence-specific nor richly searchable. Microbiome research spans a wide range of fields including ecology, epidemiology, medicine, biotechnology, and agricultural engineering, and is likely to become more integrative in the future (34). Synthesis efforts to bridge knowledge gaps across environments (6) will likely rely on the ability to find data by searching databases directly, rather than resorting to a body of literature which is currently spread across the journals from various fields. It is therefore essential that microbiome data is deposited to the appropriate INSDC databases, which also store searchable metadata, to ensure future reusability, and that current databases continue to make improvements to increase the searchability of their databases.

### Data deposition

Due to the sensitive nature of unpublished data, INSDC databases allow users to upload their data and receive an accession number but keep the data private indefinitely (35). This was evident in our data collection. Among the 2,015 articles which listed accession numbers, 7.2% (n=146) of the articles had listed accession numbers correctly but had not made the sequence data public, and this proportion increased slightly over time (χ^2^= 3.9, *p*=0.05, Supplementary Figure S2), indicating that recent articles were more likely to have not made their data public at the time of manuscript publication. Overall, the proportion of studies with non-public data rose from 5.9% in 2015 to 12.2% in 2019 (Supplementary S2). We also found that 2.2% of the studies (n=45) listed incorrect accession numbers (Supplementary Figure S2). Over the period studied, this proportion went up significantly (χ^2^ = 9.18, *p*<0.001), from 1.3% in 2015 to 5.3% in 2019.

An additional 2.5% of the studies (n=51) had not made their sequence metadata public, a trend which increased over the period studied (χ^2^ = 14.83, *p*<0.001, Supplementary S2). Among the studies targeting the V3-V4 region of the 16S rRNA gene, we found that 33 studies had deposited incorrect accession numbers over the period studied.

### Data format

While microbiome sequence data has been lauded for its uniform format, we found that the sequence files deposited varied quite widely in the format in which they were deposited, often rendering them unusable. Among the subset of 441 studies targeting the V3-V4 region of the 16S rRNA gene for which INSDC-compliant accession numbers were available and data was public in the repository (representing 45,440 samples), we found that 11.8% of the studies (n=52) had uploaded a single sequence file for the entirety of the project, despite analyzing more than one sample (Figure 2). Currently, most sequencing platforms are able to output demultiplexed data, i.e., one or more sequence file(s) per sample. However, common legacy formats consisted of one or two files for the entirety of the run as well as a mapping file, which contained the primer barcodes used to demultiplex the sequences. INSDC platforms require sequencing data to be demultiplexed prior to deposition, rendering non-demultiplexed raw data unusable due to elimination of any header information in the sequence files. Our data reflected this legacy effect: between 2015 and 2019, the proportion of studies which contained a single sequence file decreased significantly from 24.5% to 9.5% (χ^2^= 16.92, *p*<0.001, Figure 2). Furthermore, over this period, the proportion of studies which used Illumina platforms increased, and the proportion which used the older 454 pyrosequencing technique decreased (χ^2^= 10.96, *p*<0.001; and χ^2^= 10.46, *p*=0.001, respectively; Figure 2). Our findings shed light on the effect that fluxes in sequencing platform and file formats have on the scientific community’s ability to access data later.

**Figure 2.**
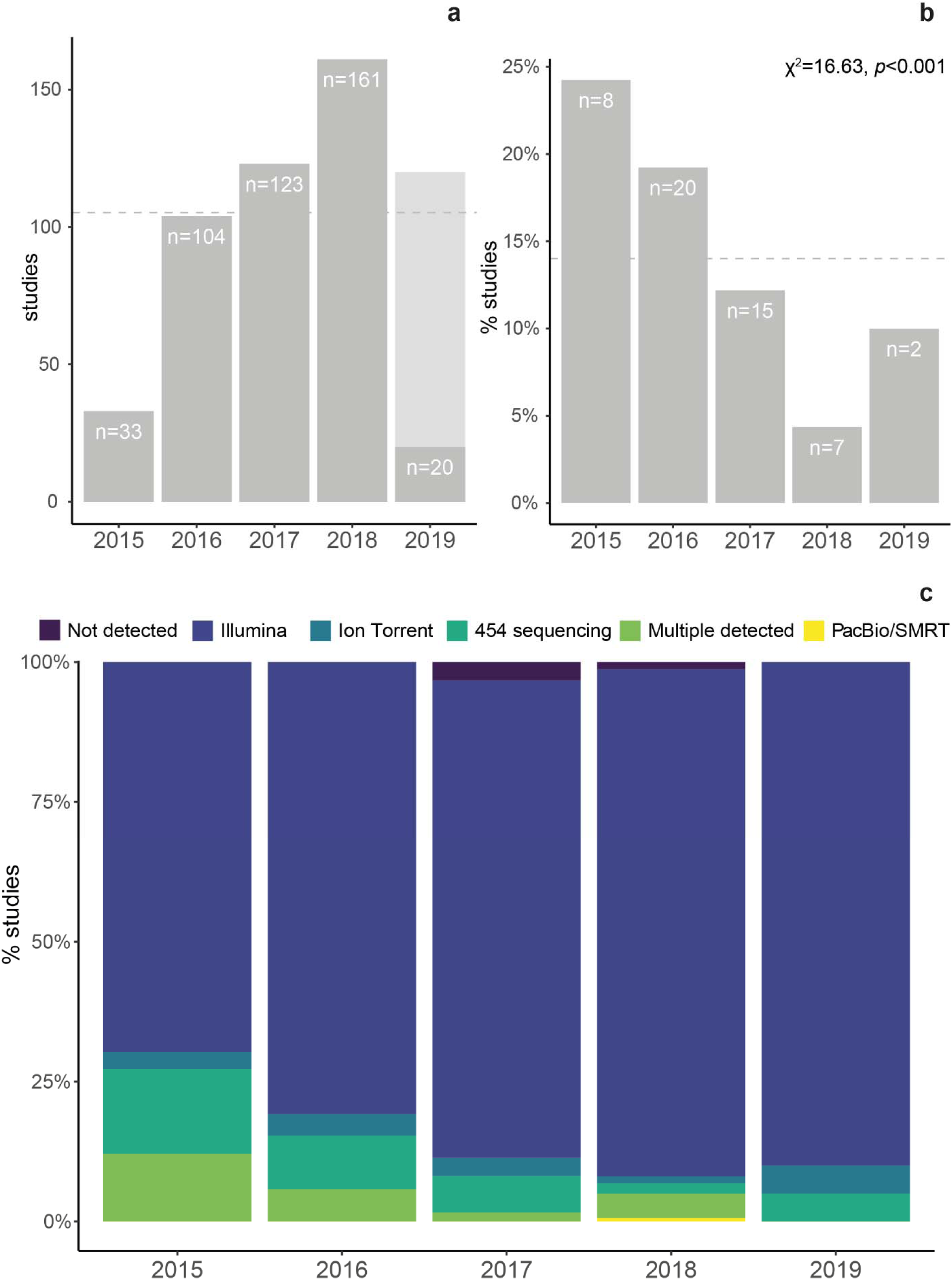
Trends in community sequencing practices over time. The number of amplicon sequencing studies targeting the V3-V4 region of the 16S rRNA gene (a). The total number of articles for 2019 was estimated from the first two months of data (light grey). The proportion of these studies which were deposited in a single sequence file, a data deposition error associated with legacy sequence formats (b) significantly decreased over the period studied (evaluated with a Chi-squared test for trend in proportions). The mean proportion of studies deposited in this manner through all years is indicated with a grey dashed line. The proportion of each sequencing platform used across the studies, over time (c). As preferences for sequencing techniques continue to change, so will the needs of users seeking to archive their data.

We found an additional 1.6% of the studies (n=7) which contained sequence files lacking standard quality scores. During sequence processing, quality scores allow users to assess the quality of the data and to exclude sequence reads with poor quality. Therefore, sequence data lacking quality scores is not reusable. We also found that 18.1% (n=80) of the studies contained putative primer sequences, but there was no significant change over time (χ^2^= 2.33, *p*=0.13, Supplementary Figure S3). Primer presence is not a strong determinant of whether data is reusable, and it is advised that data is archived in the rawest format possible. However, knowledge of primer presence and primer sequence identity are essential in the proper reprocessing of the data in the future, and currently, there are no standard methodologies for including this information in the metadata. Without this information, barcode and primer sequences may be interpreted as regular data.

### Data labeling

Properly labeling sequence data and including detailed metadata is essential to data reuse (36). Among the studies which targeted the V3-V4 region of the 16S rRNA gene and provided accession numbers, errors in labeling exceeded any other type of error (Figure 3). Because our data collection contained 16S rRNA amplicon sequencing studies exclusively, we checked whether this information was included correctly in the sequence metadata. Among these studies, 12% (n=53) of them had incorrectly labeled their sequences, using terms other than “Amplicon”. The percentage of studies with this error varied widely, from 18.2% (n=6) in 2015 to 5% (n=1) in 2019, no trends were found over time (χ^2^= 1.41, *p*=0.24, Supplementary Figure S3).

**Figure 3.**
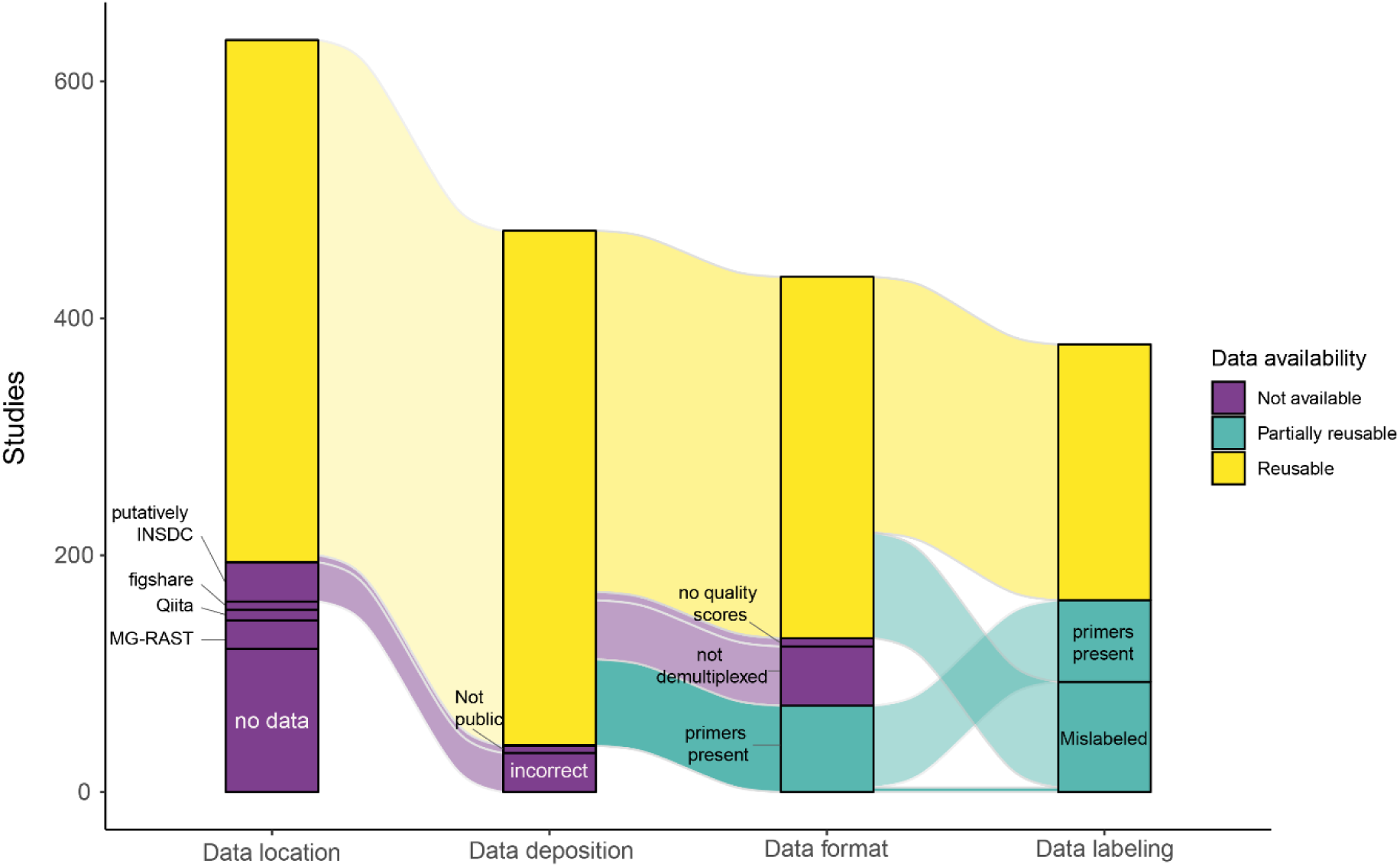
The fate of microbiome community data. An assessment of the data location and state of the 635 studies which sequenced the V3-V4 hypervariable region of the 16S rRNA gene. Data loss was divided into four categories: loss due to data location, errors in data deposition, errors in data formatting, and errors in data labeling. Data was categorized as ‘reusable’ no faults in the above four categories were found. Data was categorized as ‘partially usable’ if faults in data formatting or data labelling were likely to create obstacles in data reuse (i.e., if data not findable in the database due to mislabeling). Finally, data was categorized as ‘not available’ if it was not publicly available on INSDC databases, or if the datasets were missing data which precluded their reusability.

A defining development in the field of microbial ecology has been the advent of paired-end sequencing, by which both ends of the fragment are sequenced and later aligned *in silico*, resulting in a higher read accuracy or in longer read lengths (37). Next-generation sequencers currently output forward and reverse reads in separate files. We checked whether datasets labeled as “paired” also contained files corresponding to forward and reverse reads, or appropriately labeled reads. This was not the case for 16.8% of the studies targeting the V3-V4 region (n=74), and exhibited no temporal patterns (χ^2^= 2.09, *p*=0.15, Supplementary Figure S3). Much like datasets which include putative primer sequences, when data is labeled as paired-ended and only a single file per sample is available, future users must infer what the true state of the sequence data is. Upon a qualitative inspection of these datasets, we found that a common source of the error was that only the forward reads or merged reads had been deposited. This labeling error does not render the data unusable, but makes the sequencing conditions hard to understand for future users, who must reverse engineer the methods from the data format and quality information.

### Repopulating the archives

Errors in data deposition may render entire datasets unavailable for future research, or they may greatly complicate future data reuse. To this end, we followed the 635 studies which performed amplicon sequencing of the V3-V4 segment of the 16S rRNA gene (Figure 3). Throughout the process of archiving data, we found that 19% were not archived at all, while 6.3% of the datasets were archived in other databases which were not designed for this task. A further 6.1% datasets were improperly deposited to sequence databases, while 11.5% and 8.9% were made partially (i.e., contained putative primer sequences) or completely unavailable (i.e., not demultiplexed) due to errors in data formatting, respectively. Finally, errors in labeling affected 14.6% of the available data.

In total, only 34% of the studies identified (n=216) contained fully reusable datasets, and 25.5% (n=162) contained partially available datasets. A further 40.3% of studies (n=256) contained data that was either not available or not reusable, severely limiting advances in synthetic microbiome research and compromising some of the fundamental principles in science (12). An additional hurdle to data reuse is the availability of suitable metadata. While a detailed analysis is beyond the scope of this study, an assessment of the content and informational value of the metadata supplied for the 441 V3-V4 studies is presented in Figures S4-S6 and S8.

Our findings show the true extent of reusability of the sequencing data which has been deposited over the past 5 years, and reveal a serious gap between the sequence data which is uploaded and that which may serve to inform future research. By identifying the main reasons for data loss (i.e., loss due to data location, errors in data deposition, errors in data formatting, and errors in data labeling), the present study provides the basis of and concrete recommendations for improved data archiving practices (Table 1). Given the plethora of pressing environmental, biotechnological, and medical challenges, preserving microbiome data is particularly relevant across fields of basic and applied research.

**Table 1.** Recommendations for the future improvement of data archiving practices

| studies | Issue | Recommendations |
| --- | --- | --- |

| affected |  |  |
| --- | --- | --- |
| 31.4 % | Data is not accessible<br>• Data is not deposited<br>• Data is not deposited to INSDC-affiliated databases<br>• Accession numbers are incorrect<br>• Data is private<br>• Metadata is private | **Researchers:**<br>• Make deposited data available upon a manuscript's publication<br>• Ensure accession numbers are correct in the published article.<br>**Publishers:**<br>• Require that the sequencing data is available upon article submission, and remind authors to make the data publicly available by the time of publication (38).<br>• Demand that datasets are deposited to the appropriate INSDC databases prior to submission in order to guarantee their long-term availability.<br>**Data archives:**<br>• Require that users select a date to make data public during the deposition process. |
| 23.6% | Changes in data formatting practices<br>• Data is uploaded in legacy file formats<br>• Single sequence files are uploaded for paired-end data | **Researchers:**<br>• Ensure that a minimum set of data is provided in order to allow for reproducibility. This includes formally collecting and depositing metadata to include experiment, sample, and sequence information; and recording protocols using modern tools (39) (i.e., protocols.io for laboratory protocols and RNotebook or Jupyter Notebook for bioinformatics code)<br>**Data archives:**<br>• Allow for the deposition of more diverse sequence file types, (i.e., allow for the deposition of mapping files)<br>• Develop new standards which require the reporting of sequencing processing metadata. Essential information such as DNA extraction, sequencing, and processing protocols should be providable via a DOI.<br>• Have a common and precise language regarding 'best practices' for data deposition (e.g., the inclusion of primers)(17).<br>• Keep publicly available changelogs of database guidelines, so that users may understand how and why data was deposited in a particular format in the past. |
| 14.6% | Mislabeling<br>• Amplicon sequences not listed as 'amplicon'<br>• Single sequence files are uploaded for paired-end data | **Researchers:**<br>• Become familiarized with the terms associated with sequencing and sequence formats for proper data upload (40).<br>• Proactive interaction with database holders (i.e., helpdesk) to ensure that data deposition is done correctly.<br>**Publishers:**<br>• Demand that the metadata tables be included during article submission for peer review |
|  |  | **Data archives:**<br>Recognize that amplicon sequencing is an increasingly interdisciplinary technique, and continue the current trend towards improved documentation and explanations. In particular, users may benefit from more precise guidelines into what constitutes informative metadata for the purposes of archiving (e.g., listing the environment as ‘human’ vs. ‘human gut’, Figure S4) |

## Data availability statement

The data that support the findings of this study are available as supplementary material. Note that variables identifying specific articles (i.e., authors, DOI) will be removed prior to publication. The pattern-based text extraction algorithms are available at https://github.com/komax/teitocsv. The R code used for data analysis is available to reviewers as supplementary material and will become publicly available in the authors’ GitHub accounts upon acceptance.

## Acknowledgements

We would like to thank H.R.P. Phillips for help with data visualization, and J. Chase, R. van Klink and S. Tem for the valuable discussions. The authors declare no conflict of interests.

## Author contributions

SDJ conceived of the study, wrote the manuscript and created the figures. MK designed and built the pipeline to extract textual patterns from the PDFs on the entire corpus. AHB performed all bioinformatics analyses of the sequence data in our collection. NE contributed to the organization and presentation of the results. All authors contributed substantially to subsequent revisions.

## References

1. Steen, A.D., Crits-Christoph, A., Carini, P., DeAngelis, K.M., Fierer, N., Lloyd, K.G. and Cameron Thrash, J. (2019) High proportions of bacteria and archaea across most biomes remain uncultured. ISME J., 10.1038/s41396-019-0484-y.

2. Lloyd, K.G., Steen, A.D., Ladau, J., Yin, J. and Crosby, L. (2018) Phylogenetically Novel Uncultured Microbial Cells Dominate Earth Microbiomes. mSystems, 3, 1–12.

3. Kodama, Y., Shumway, M. and Leinonen, R. (2012) The sequence read archive: Explosive growth of sequencing data. Nucleic Acids Res., 40, 2011–2013.

4. Locey, K.J. and Lennon, J.T. (2016) Scaling laws predict global microbial diversity. Proc. Natl. Acad. Sci., 113, 5970 LP – 5975.

5. Shade, A., Dunn, R.R., Blowes, S.A., Keil, P., Bohannan, B.J.M.M., Herrmann, M., Küsel, K., Lennon, J.T., Sanders, N.J., Storch, D., et al. (2018) Macroecology to Unite All Life, Large and Small. Trends Ecol. Evol., 33, 731–744.

6. Langenheder, S. and Lindström, E.S. (2019) Factors influencing aquatic and terrestrial bacterial community assembly. Environ. Microbiol. Rep., 11, 306–315.

7. Stegen, J.C., Bottos, E.M. and Jansson, J.K. (2018) A unified conceptual framework for prediction and control of microbiomes. Curr. Opin. Microbiol., 44, 20–27.

8. Thompson, L.R., Sanders, J.G., McDonald, D., Amir, A., Ladau, J., Locey, K.J., Prill, R.J., Tripathi, A., Gibbons, S.M., Ackermann, G., et al. (2017) A communal catalogue reveals Earth’s multiscale microbial diversity. Nature.

9. Wirbel, J., Pyl, P.T., Kartal, E., Zych, K., Kashani, A., Milanese, A., Fleck, J.S., Voigt, A.Y., Palleja, A. and Ponnudurai, R. (2019) Meta-analysis of fecal metagenomes reveals global microbial signatures that are specific for colorectal cancer. Nat. Med., 25, 679.

10. Parks, D.H., Rinke, C., Chuvochina, M., Chaumeil, P.-A., Woodcroft, B.J., Evans, P.N., Hugenholtz, P. and Tyson, G.W. (2017) Recovery of nearly 8,000 metagenome-assembled genomes substantially expands the tree of life. Nat. Microbiol., 2, 1533.

11. Rocca, J.D., Simonin, M., Blaszczak, J.R., Ernakovich, J.G., Gibbons, S.M., Midani, F.S. and Washburne, A.D. (2019) The Microbiome Stress Project: Toward a global meta-analysis of environmental stressors and their effects on microbial communities. Front. Microbiol., 10.

12. Wilkinson, M.D., Dumontier, M., Aalbersberg, Ij.J., Appleton, G., Axton, M., Baak, A., Blomberg, N., Boiten, J.-W., da Silva Santos, L.B. and Bourne, P.E. (2016) The FAIR Guiding Principles for scientific data management and stewardship. Sci. data, 3.

13. Hampton, S.E., Strasser, C.A., Tewksbury, J.J., Gram, W.K., Budden, A.E., Batcheller, A.L., Duke, C.S. and Porter, J.H. (2013) Big data and the future of ecology. Front. Ecol. Environ., 11, 156–162.

14. Roche, D.G., Kruuk, L.E.B., Lanfear, R. and Binning, S.A. (2015) Public Data Archiving in Ecology and Evolution□: How Well Are We Doing□? 10.1371/journal.pbio.1002295.

15. Karsch-Mizrachi, I., Nakamura, Y. and Cochrane, G. (2012) The International Nucleotide Sequence Database Collaboration. 40, 33–37.

16. Zhou, Z., Wang, C. and Luo, Y. (2018) Effects of forest degradation on microbial communities and soil carbon cycling: A global meta□analysis. Glob. Ecol. Biogeogr., 27, 110–124.

17. Yilmaz, P., Kottmann, R., Field, D., Knight, R., Cole, J.R., Amaral-Zettler, L., Gilbert, J.A., Karsch-Mizrachi, I., Johnston, A., Cochrane, G., et al. (2011) Minimum information about a marker gene sequence (MIMARKS) and minimum information about any (x) sequence (MIxS) specifications. Nat. Biotechnol., 29, 415–420.

18. Gilbert, J.A., Jansson, J.K. and Knight, R. (2014) The Earth Microbiome project: successes and aspirations. BMC Biol., 12, 69.

19. Harzing, A.W. (2007) Publish or Perish.

20. Schloss, P.D., Westcott, S.L., Ryabin, T., Hall, J.R., Hartmann, M., Hollister, E.B., Lesniewski, R.A., Oakley, B.B., Parks, D.H. and Robinson, C.J. (2009) Introducing mothur: open-source, platform-independent, community-supported software for describing and comparing microbial communities. Appl. Environ. Microbiol., 75, 7537–7541.

21. Caporaso, J.G., Kuczynski, J., Stombaugh, J., Bittinger, K., Bushman, F.D., Costello, E.K., Fierer, N., Pena, A.G., Goodrich, J.K. and Gordon, J.I. (2010) QIIME allows analysis of highthroughput community sequencing data. Nat. Methods, 7, 335–336.

22. Bolyen, E., Rideout, J.R., Dillon, M.R., Bokulich, N.A., Abnet, C., Al-Ghalith, G.A., Alexander, H., Alm, E.J., Arumugam, M. and Asnicar, F. (2018) QIIME 2: Reproducible, interactive, scalable, and extensible microbiome data science PeerJ Preprints.

23. Caporaso, J.G., Lauber, C.L., Walters, W.A., Berg-Lyons, D., Huntley, J., Fierer, N., Owens, S.M., Betley, J., Fraser, L. and Bauer, M. (2012) Ultra-high-throughput microbial community analysis on the Illumina HiSeq and MiSeq platforms. ISME J., 6, 1621–1624.

24. Sayers, E.W., Agarwala, R., Bolton, E.E., Brister, J.R., Canese, K., Clark, K., Connor, R., Fiorini, N., Funk, K. and Hefferon, T. (2019) Database resources of the national center for biotechnology information. Nucleic Acids Res., 47, D23.

25. Martin, M. (2011) Cutadapt removes adapter sequences from high-throughput sequencing reads. EMBnet. J., 17, 10–12.

26. Apprill, A., McNally, S., Parsons, R. and Weber, L. (2015) Minor revision to V4 region SSU rRNA 806R gene primer greatly increases detection of SAR11 bacterioplankton. Aquat. Microb. Ecol., 75, 129–137.

27. Parada, A.E., Needham, D.M. and Fuhrman, J.A. (2016) Every base matters: assessing small subunit rRNA primers for marine microbiomes with mock communities, time series and global field samples. Environ. Microbiol., 18, 1403–1414.

28. Andrews, S. (2010) FastQC: a quality control tool for high throughput sequence data.

29. Kans, J. (2020) Entrez direct: E-utilities on the UNIX command line. In *Entrez Programming Utilities Help [Internet]*. National Center for Biotechnology Information (US).

30. Team, R.C. (2017) R: A language and environment for statistical computing. R Foundation for Statistical Computing, Vienna, Austria. 2016.

31. Gonzalez, A., Navas-Molina, J.A., Kosciolek, T., McDonald, D., Vázquez-Baeza, Y., Ackermann, G., DeReus, J., Janssen, S., Swafford, A.D. and Orchanian, S.B. (2018) Qiita: rapid, web-enabled microbiome meta-analysis. Nat. Methods, 15, 796.

32. Keegan, K.P., Glass, E.M. and Meyer, F. (2016) MG-RAST, a metagenomics service for analysis of microbial community structure and function. In Microbial Environmental Genomics (MEG). Springer, pp. 207–233.

33. Caporaso, J.G., Lauber, C.L., Walters, W.A., Berg-Lyons, D., Lozupone, C.A., Turnbaugh, P.J., Fierer, N. and Knight, R. (2011) Global patterns of 16S rRNA diversity at a depth of millions of sequences per sample. Proc. Natl. Acad. Sci., 108, 4516 LP – 4522.

34. Craven, D., Winter, M., Hotzel, K., Gaikwad, J., Eisenhauer, N., Hohmuth, M., König□Ries, B. and Wirth, C. (2019) Evolution of interdisciplinarity in biodiversity science. Ecol. Evol.

35. National Center for Biotechnology Information (2010) SRA Handbook.

36. Lima, M.S. and Smith, D.R. (2017) Don’t just dump your data and run. 18, 2087–2089.

37. Bartram, A.K., Lynch, M.D.J., Stearns, J.C., Moreno-Hagelsieb, G. and Neufeld, J.D. (2011) Generation of multimillion-sequence 16S rRNA gene libraries from complex microbial communities by assembling paired-end Illumina reads. Appl. Environ. Microbiol., 77, 3846–3852.

38. Vines, T.H., Andrew, R.L., Bock, D.G., Franklin, M.T., Gilbert, K.J., Kane, N.C., Moore, J.-S., Moyers, B.T., Renaut, S. and Rennison, D.J. (2013) Mandated data archiving greatly improves access to research data. FASEB J., 27, 1304–1308.

39. Rambold, G., Yilmaz, P., Harjes, J., Klaster, S., Sanz, V., Link, A., Glöckner, F.O. and Triebel, D. (2019) Meta-omics data and collection objects (MOD-CO): a conceptual schema and data model for processing sample data in meta-omics research. Database, 2019.

40. Marchesi, J.R. and Ravel, J. (2015) The vocabulary of microbiome research: a proposal. Microbiome, 3, 31.

