## Supplementary for "The archives are half-empty: a field-wide assessment of the availability of microbial community sequencing data"

**Supplementary materials**

Table S1. Journals included in this study. We selected 17 of the most prominent journals for the publication of bacterial community, or microbiome research, and retrieved all of their publications since 2015. We screened 26,927 articles and selected the 2015 which mentioned the 16S rRNA gene, and contained an INSDC-compliant accession number. For more in-depth analyses, we sub-selected the 441 articles which had sequenced the V3-V4 hypervariable region of the 16S rRNA gene, between base pairs 515 and 806. We found 33 cases in which putative accession numbers were detected but did not match any INSDC database's public records, and were excluded from further investigation.

| Journal name | 16S rRNA sequencing studies | 16S rRNA sequencing studies, V3-V4 region |
|---|---|---|
| Annals of Microbiology | 17 | 5 |
| Applied and Environmental Microbiology | 207 | 51 |
| BMC Genomics | 15 | 3 |
| BMC Microbiology | 63 | 14 |
| eLife | 5 | 1 |
| Environmental Microbiology | 203 | 42 |
| Environmental Microbiology Reports | 32 | 6 |
| FEMS Microbiology Ecology | 189 | 40 |
| Frontiers in Microbiology | 854 | 192 |
| ISME Journal | 118 | 18 |
| Journal of Applied Microbiology | 41 | 9 |
| Journal of Microbiology | 5 | 1 |
| Journal of Microbiological Methods | 2 | 1 |
| Journal of Microbiology Korea | 9 | 2 |
| mBio | 40 | 7 |
| BMC Microbiome | 204 | 47 |
| Nature Microbiology | 11 | 2 |

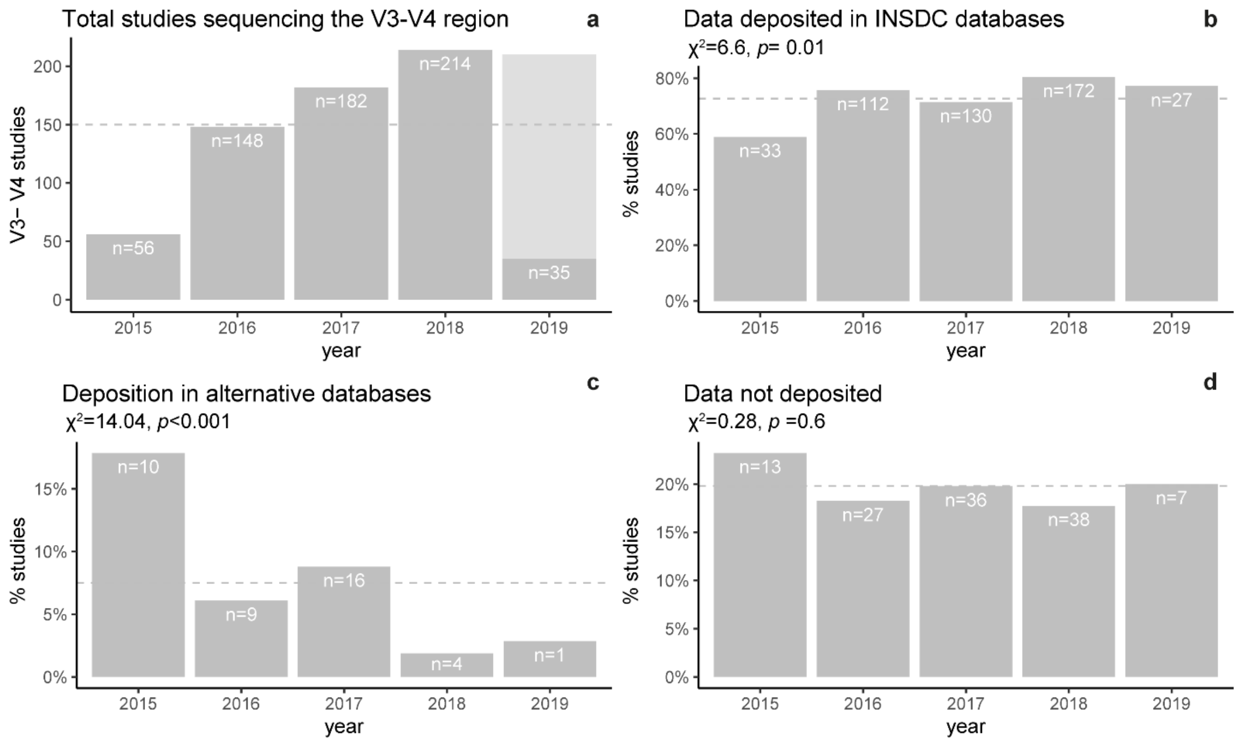


Figure S1. Increasing popularity of INSDC databases. The number of studies targeting the V3-V4 region of the 16S rRNA grew over the period studied (a), as did the proportion of these studies which was deposited to INSDC databases (b); however this increase was driven by an increased preference for INSDC databases among existing databases, as evidenced by the decrease in studies depositing their data in alternative databases (c) and the constant proportion of studies for which we did not detect any data deposition (d). The total number of articles for 2019 was estimated from the first two months of data (light grey). For panels b-d, grey dashed lines indicate the mean percentage of studies which fulfilled each condition over the period studied. Trends over time were evaluated with a Chi-squared test for trend in proportions.

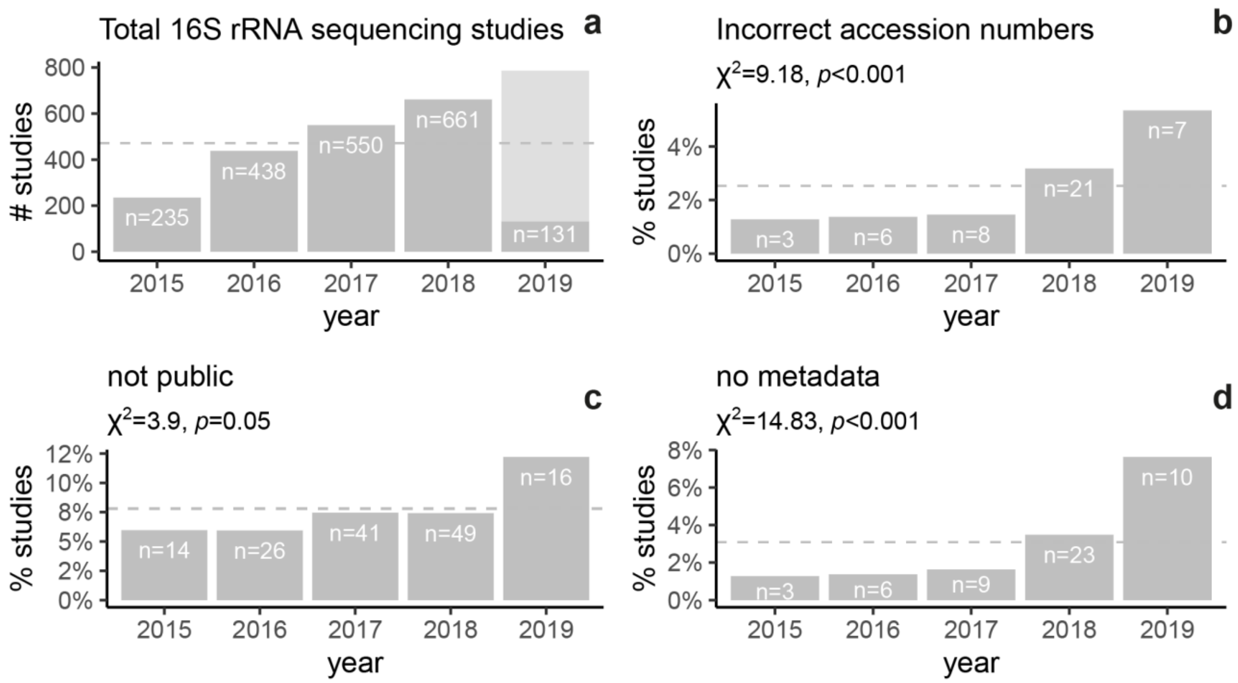


Figure S2. Increase in data loss due to errors in data deposition over time. Over the period studied, the number of articles sequencing the 16S rRNA gene and depositing the sequences to INSDC databases grew (a), as did the proportion of studies which listed incorrect accession numbers (b), kept their data private after the article's acceptance (c), or did not submit metadata (d). The total number of articles for 2019 was estimated from the first two months of data (light grey). For panels b-d, grey dashed lines indicate the mean percentage of studies which fulfilled each condition over the period studied. Trends over time were evaluated with a Chi-squared test for trend in proportions.

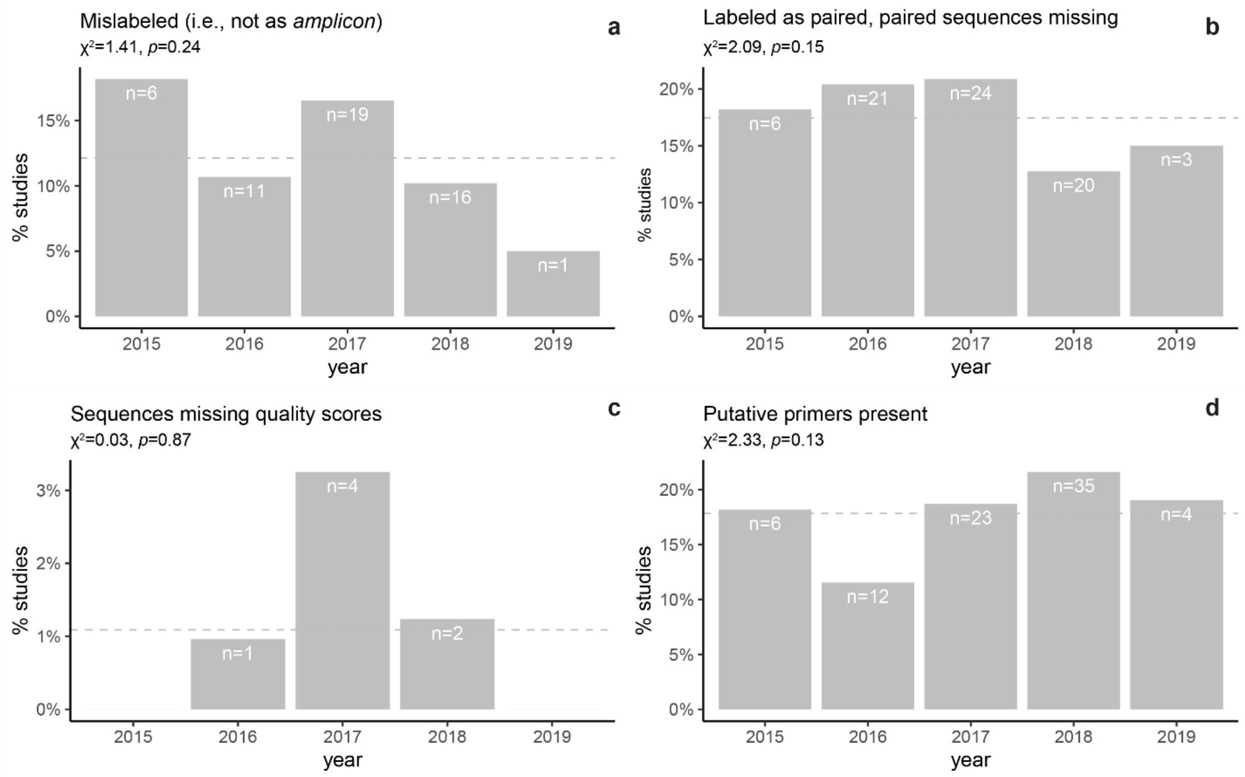


Figure S3. Errors in metadata labeling and data formatting over the period studied. Datasets labeled as containing sequence types other than 'amplicon' (a) and datasets labeled as paired-ended but lacking paired sequence files (b) were counted, as well as the number of studies which supplied sequence data lacking quality scores (c), or including putative primer sequences (d). Grey dashed lines indicate the mean percentage of studies which fulfilled each condition over the period studied. Trends over time were evaluated with a Chi-squared test for trend in proportions. Proportions were calculated relative to the studies targeting the V3-V4 region of the 16S rRNA gene which deposited their data in INSDC databases.

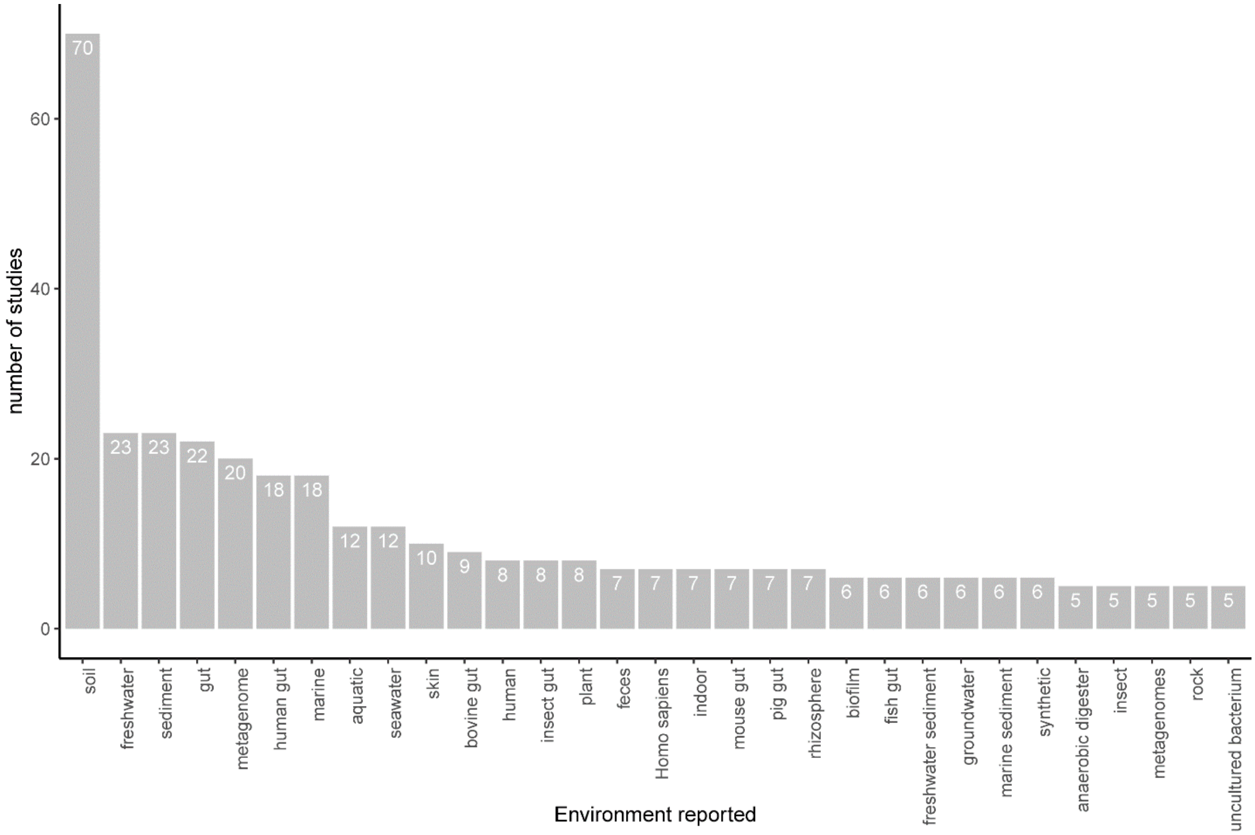


Figure S4. Environments targeted by studies sequencing the V3-V4 region of the 16S rRNA gene. We selected the 441 V3-V4 studies for which sequence data and metadata was available. Environment metadata was extracted from the 'ScientificName' attribute of the metadata files obtained from NCBI. Only environments which were listed in at least 5 studies are included. Among the 441 studies, 172 distinct environments were reported. Note that 25 studies listed the environment as 'metagenome(s)', and different degrees of ambiguity exist in listing the environment (i.e., gut and *Homo sapiens* vs. human gut).

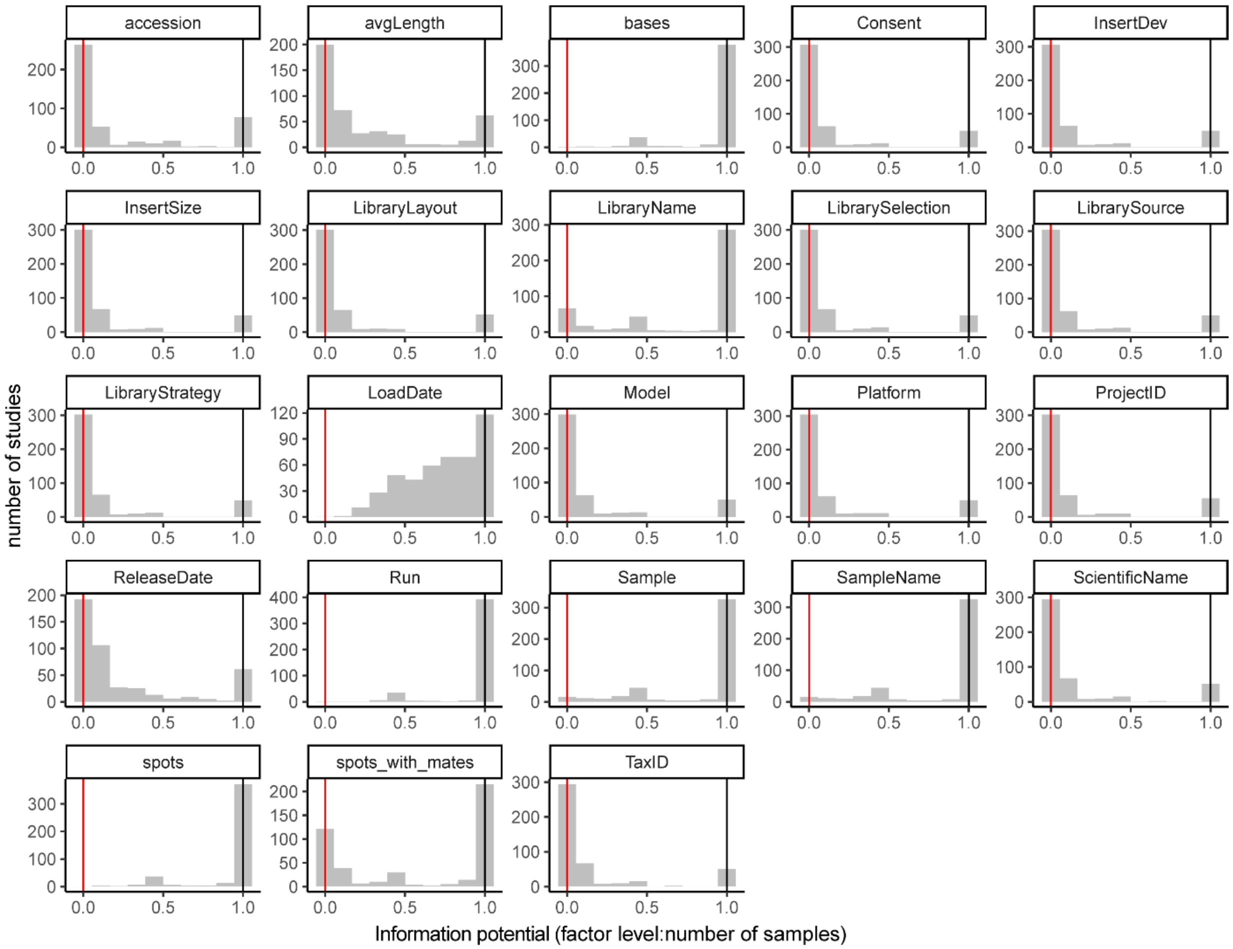


Figure S5. The informative potential of mandatory metadata fields and tax ID. Metadata was assessed for the 441 studies targeting the V3-V4 hypervariable region of the 16S rRNA gene for which metadata was available. Fields presented here are primarily technical, and include metadata on sequencing parameters (i.e, InsertSize) as well as experimental information (i.e., SampleName). If a field is the same for the entire study, it is tallied on the left end of the distribution (red line). If the field is unique for each sample in the experiment, the number of factor levels and samples is the same, and it is tallied on the right side of the distribution (black line). From an experimental perspective the most informative fields are those which vary but occur multiple times within the experiment. These would appear in the center of the distribution, as is the case with the fields ‘LibraryName’ and ‘sample’.

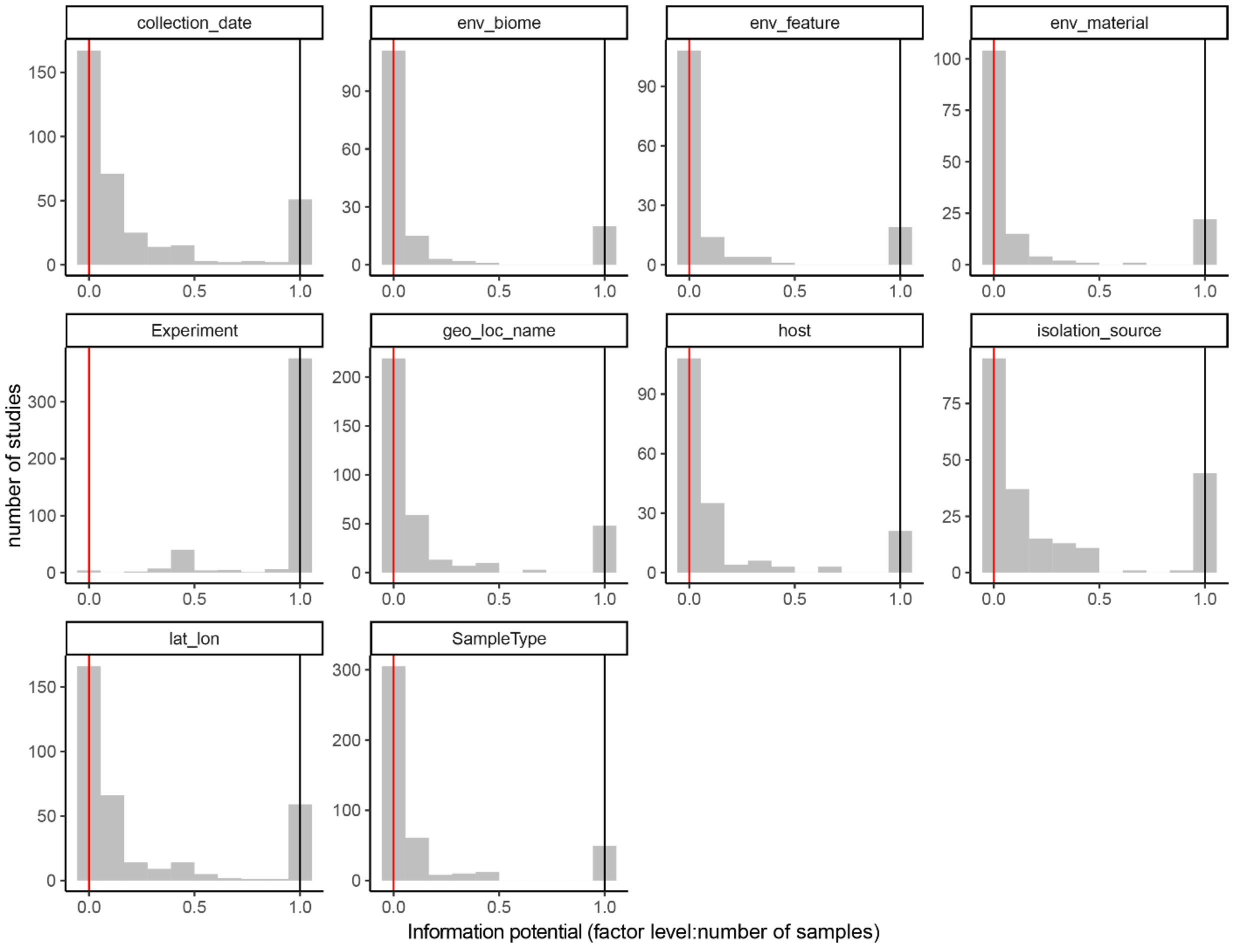


Figure S6. The informative potential of optional, popular metadata fields. Metadata was assessed for the 441 studies targeting the V3-V4 hypervariable region of the 16S rRNA gene for which metadata was available. Popular sample metadata fields were considered those for which data was available in at least 111 datasets (25%). Note that whether a field is optional or mandatory may change over time as INSDC deposition policies are improved. Fields presented here include experimental information (i.e., SampleType). From an experimental perspective the most informative fields are those which vary but occur multiple times within the experiment. These would appear in the center of the distribution,

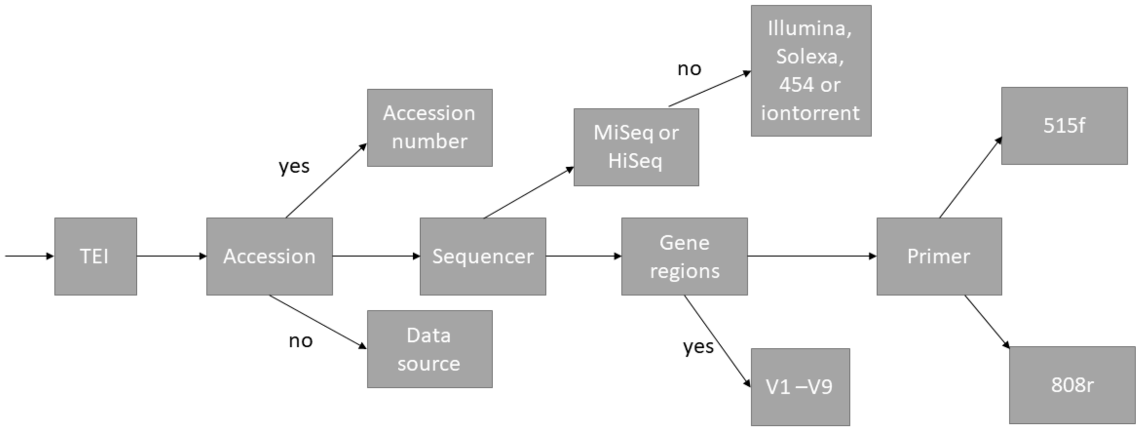


Figure S7. Pipeline to process TEI XML documents in order to look up accession numbers, sequencing techniques, gene regions and primers in the paper's title or otherwise the main text.

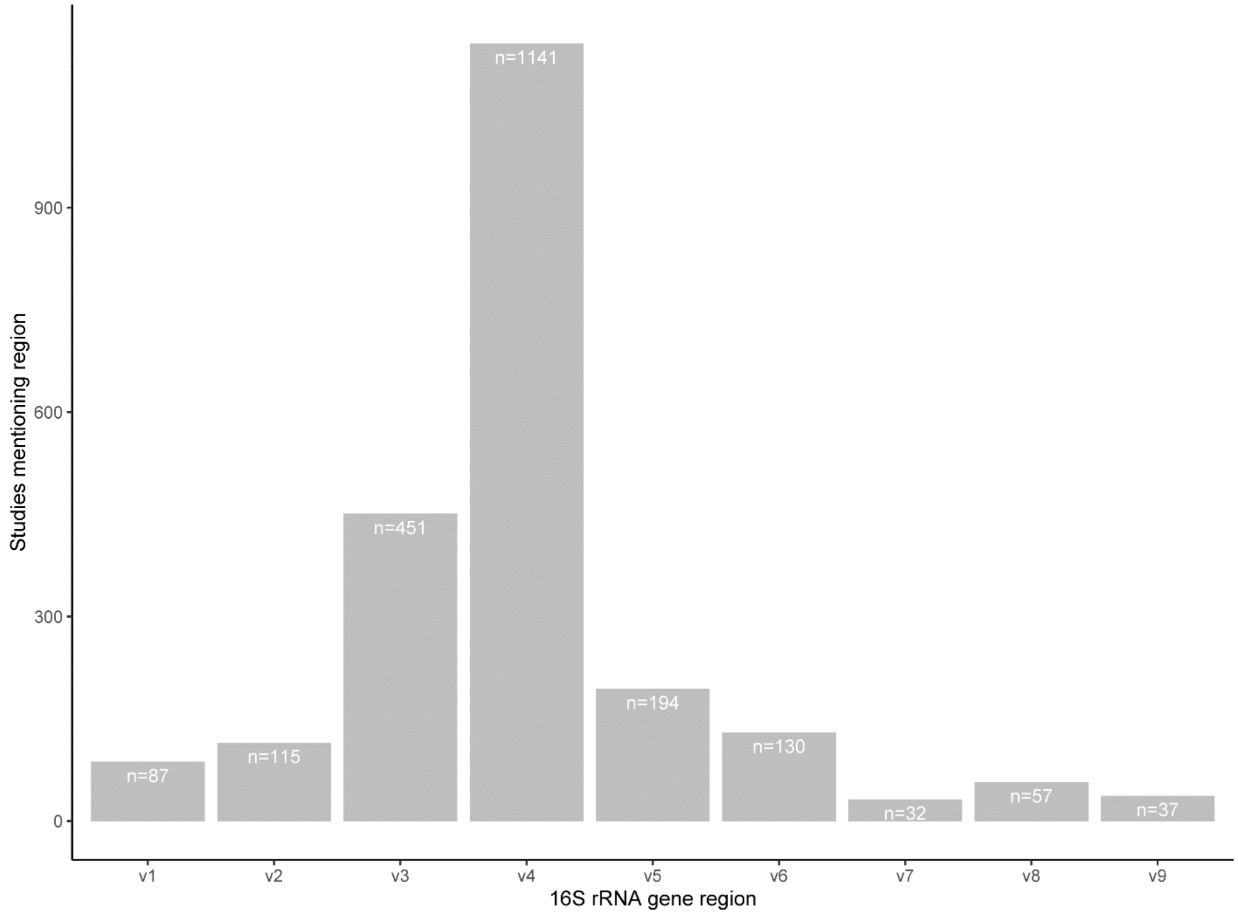


Figure S8. Frequency of sequencing for each of the 16S rRNA hypervariable regions. For this assessment, we considered 2245 studies in our database which contained the keyword “16S” and mentioned at least one hypervariable region. When more than one region was mentioned, both were considered. Studies targeting the V3-V4 hypervariable region were selected for further inspection.
